# Chronic optogenetic stimulation of hippocampal engrams variably modulates social behaviors in mice

**DOI:** 10.1101/2020.05.28.121822

**Authors:** Emily Doucette, Heloise Leblanc, Amy Monasterio, Christine Cincotta, Stephanie L. Grella, Jesse Logan, Steve Ramirez

## Abstract

The hippocampus processes both spatial-temporal information and emotionally salient experiences. To test the functional properties of discrete sets of cells in the dorsal dentate gyrus (dDG), we examined whether chronic optogenetic reactivation of these ensembles was sufficient to modulate social behaviors in mice. We found that chronic reactivation of dDG cells in male mice was sufficient to enhance social behaviors in a female exposure task when compared to pre-stimulation levels. However, chronic reactivation of these cells was not sufficient to modulate group differences in a separate subset of social behaviors, and multi-region analysis of neural activity did not yield detectable differences in immediate-early gene expression or neurogenesis, suggesting a dissociation between our chronic stimulation-induced behavioral effects and underlying neural responses. Together, our results demonstrate that chronic optogenetic stimulation of cells processing valent experiences enduringly and unidirectionally modulates social interactions between male and female mice.

## 1. Introduction

Social behaviors are dramatically impaired across many psychiatric disorders, though the underlying mechanisms sufficient to precipitate or alleviate such impairments remain largely unknown. Promisingly, previous studies have demonstrated that chronic optogenetic reactivation of both cell bodies and projection-specific elements can “reprogram” circuit-level and behavioral outputs in healthy and maladaptive states (Creed, Pascoli, & Lüscher, 2019; Tye, 2014). However, the behavioral effects of chronic optogenetic stimulation of memory ensembles are region-specific and experience dependent. Specifically, reactivating dDG cells that were active during a positive experience was sufficient to rescue depressive-like behavior in mice, while chronically reactivating dDG cells previously active during fear conditioning was sufficient to lastingly suppress or enhance a context-specific memory (Ramirez et al., 2015; Chen et al., 2019). To test whether or not our chronic stimulation strategy generalizes to other behaviors, here we examined whether chronic optogenetic reactivation of ensembles in the dDG which are active during putative positive or negative experiences is sufficient to alter social behaviors as well as the activity of multiple brain regions.

## 2. Methods

### 2.1 Subjects

Wild-type C57BL/6N male mice (40-41 days; Charles River Laboratories) were housed with littermates in groups of 2-5 mice per cage. Mice were acclimated to the animal facility for 72 hours upon delivery before experimental procedures began and kept on a 12:12-hour light cycle (lights on at 7:00). Food and water were available *ad libitum*. Animals were put on a diet containing 40 mg/kg doxycycline (dox) after the acclimation period and 24-48 hours before receiving surgery between 6-7 weeks of age. Following surgery, mice were group-housed with littermates and were left for 10 days to recover with food and water *ad libitum* prior to experimentation. Animals were handled for 2-4 days (2 minutes per animal) at the end of the recovery period. They were also habituated to optogenetic stimulation conditions by plugging the patch cord into their headcaps and allowing them to walk around freely for 2 minutes per day for 2 days, prior to the start of the experimental period. All procedures related to mouse care and treatment were in accordance with Boston University and National Institutes of Health guidelines for the Care and Use of Laboratory animals.

### 2.2 Virus constructs and packaging; stereotaxic surgery

The pAAV_9_-*c-Fos*-tTA and pAAV_9_-TRE-ChR2-eYFP plasmids were constructed as described previously (Ramirez et al., 2013). Using these plasmids, AAV_9_ viruses were generated at the Gene Therapy Center and Vector Core at the University of Massachusetts Medical School. Viral titres were 1 × 10^13^ genome copy per milliliter for AAV_9_-TRE-ChR2-eYFP and 1.5 × 10^13^ genome copy per milliliter for AAV_9_-*c-Fos*-tTA.

All surgeries were performed under stereotaxic guidance and the following coordinates are given relative to bregma. Anesthesia was induced using 3.5% isoflurane inhalation and maintained throughout surgery at 1.5 - 2.0%. Animals received bilateral craniotomies using a 0.6 mm diameter drill bit for dDG injections. The needle was slowly lowered to the target site of - 2.2 mm AP, ±1.3 mm ML, −2.0 mm DV (relative to Bregma). A cocktail consisting of 300 nL of AAV_9_-c-Fos-tTa (300nL) + AAV_9_-TRE-ChR2-eYFP (300nL) was infused into the dDG (100nL min^−1^) using a 33-gauge needle attached to a mineral-oil filled 10 μL gastight syringe (Hamilton, #7653-01) (Figure 1A). The needle remained at the target site for 2 minutes post-injection before being slowly withdrawn. A bilateral optical fiber implant (200 μm core diameter; Doric Lenses) was lowered above the dDG injection site at −1.6 mm DV. Two bone anchor screws were secured into the skull at the anterior edges of the surgical site to anchor the implant. Layers of adhesive cement (C&B Metabond) followed by dental cement (Stoelting) were spread over the surgical site to secure the optical fiber implant to the skull. Mice received 0.1 mL buprenorphine (0.03 mg/mL i.p) and were transferred to recovery cages atop heating pads until recovery from anaesthesia. Mice were given 10 days to recover before start of experiment. Injections were verified histologically. Only data from mice with bilateral opsin expression present in the dDG were used for analyses.

**Figure 1.**
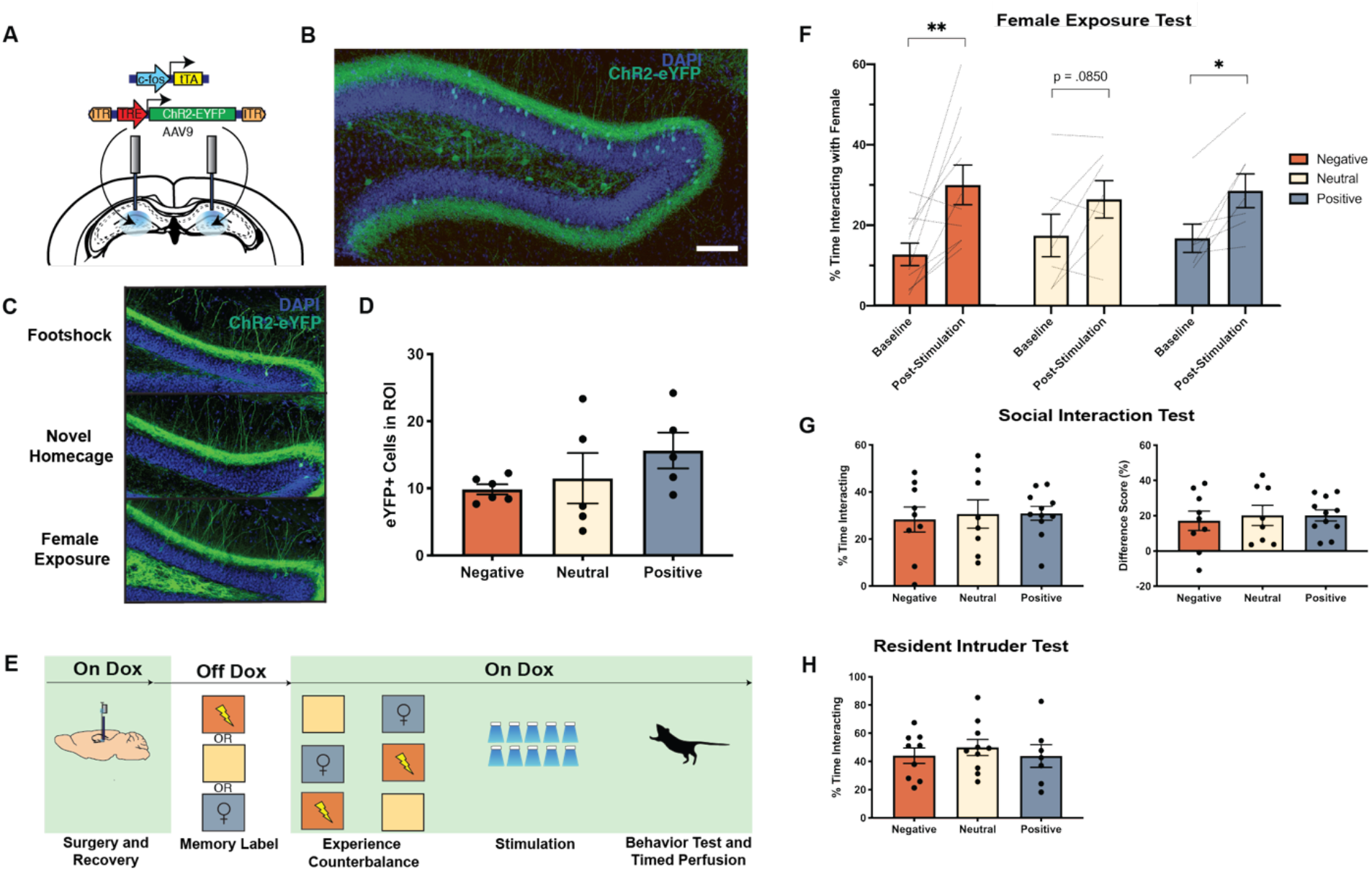
Chronic optogenetic stimulation of dDG Ensembles differentially modulates social behaviors. (**a**) Viral constructs for doxycycline (dox)-gated activity-dependent expression of ChR2 in the dDG. The immediate early gene *c-Fos* drives tetracycline transactivator (tTA), which binds to its response element (TRE) to in turn drive expression of ChR2 in a dox-regulated manner. (**b**) Representative image depicting expression of ChR2-eYFP (green) in the dDG. Scale bar represents 100 μm. **(c)** Representative images of ChR2-EYFP in DG for each group. **(d)** Ensemble sizes are not significantly different for different behavioral epochs (One-Way ANOVA, F_2,13_ = 1.392, p = 0.2834 (Negative n=6, Neutral n=5, Positive, n=5) (**d**) Behavioral schedule and groups used. Green regions depict periods in which dox was present in the diet, and white regions depict regions where dox was removed to tag active cells (“memory label”). The orange box with a shock symbol represents a four-shock protocol, the tan-colored box represents a clean homecage exposure, and the gray box with a female symbol represents exposure to a female conspecific. (**f**) Chronic stimulation of negative, neutral or positive ensembles increases female interaction time (2 Way RM ANOVA, Main Effect of Time: F _1, 21_ = 22.30, p=.0001, Post-Hoc paired t-tests Pre vs Post-stimulation: Negative p=.0073, Neutral p=.0850, Positive p=.0128 (Negative n=10, Neutral n=7, Positive n=7) (**g**) Chronically stimulating dDG ensembles encoding a negative, neutral or positive experience does not modulate post-stimulation behavior in the social interaction test or (**h**) resident intruder test. Two separate cohorts of mice underwent the protocol in a) with either the social interaction test or resident intruder test on the final day. Difference score represents the difference in time spent interacting with the conspecific cup and the empty cup in the social interaction test. Social interaction test, One-Way ANOVA Time interacting: F_2,25_ = 0.09415, p=0.9105; Difference Score: F_2,25_ = 0.1382, p = 0.8716 (Negative n=9, Neutral n=8, Positive n=11), Resident intruder test: One Way ANOVA F(2,23)=0.3150, p=0.7329 (Negative n=9, Neutral n=10, Positive n=7). Data are presented as mean ± s.e.m.

### 2.3 Neuronal tagging of behavioral epochs and counterbalanced behavior

In order to label the dDG cells active during a behavioral epoch, the dox diet is substituted with normal mouse chow; this occurs 48 hours prior to the epoch to allow for complete clearance of dox and to open the window for activity-dependent neuronal tagging (Garner et al., 2012; Liu et al., 2012) (Figure 1B-C). The mice were divided into 3 groups, and each group received a different “tagged” behavioral epoch to start, however, all mice received all behavioral epochs counterbalanced using a balanced Latin square design over a period of 3 days (negative-neutral-positive; positive-negative-neutral; neutral-negative-positive). 1) *Footshock (negative)*: animals were placed in a fear conditioning chamber and given a 4-shock protocol over a period of 500s (1.5 mA, 2s duration, 198s, 278s, 358s, 438s). 2) *Female exposure (positive)*: one female mouse (PD 30-40) was placed in a clean homecage with a clear, ventilated acrylic top. The experimental animal was then placed into the cage and allowed to freely interact with the female for 1 hour. 3) Clean homecage (neutral): mice were individually placed in a clean homecage with a clear, ventilated acrylic cage top for 500s. Immediately following the tagged behavioral epoch, mice were placed back into their homecage and again given access to dox to close the neuronal tagging window. Mice were weighed daily and monitored for health.

### 2.4 Pre-stimulation female exposure

The total amount of time that male mice interacted with a female mouse - defined as sniffing, chasing, mounting, or other contact initiated by the male - within the first 5 minutes of the 1 hour, pre-optogenetic stimulation exposure was manually scored (termed “baseline” time point).

### 2.5 Chronic optogenetic stimulation protocol

Optical stimulation was administered twice daily during the light cycle at approximately 10:00 and again at 15:00 daily for 5 days, to animals at 8-9 weeks of age. Prior to the start of the session, laser output was tested to ensure that at least 10 mW of power was delivered at the end of the patch cord (Doric Lenses). Each stimulation session lasted for 10 minutes (450 nm, 20 Hz, 15 ms pulse width) and was conducted in an almond-scented custom-built acrylic rectangular chamber with striped walls under dim lighting. The first round of behavioral tests began one day after the cessation of this protocol.

### 2.6 Post-stimulation behavioral assays (resident intruder test, social interaction test, female exposure test)

Behavioral experiments were conducted 24 hours after the final chronic optogenetic stimulation session. All behavioral assays were recorded using a web-camera (Logitech HD).

#### Resident intruder test

Experimental animals and their homecage enrichment were transferred from their homecage to a clean holding cage with their cagemates. The homecage with bedding was used as an experimental chamber with a clear, ventilated acrylic cage top. One experimental mouse from the cage was placed back into the homecage and allowed to acclimate for 1 minute, after which a novel conspecific juvenile male (PD 24-28) was introduced into the experimental male’s homecage for a 5 minute test session. Interaction was manually scored by the experimenter and was measured as experimental male-initiated behavior (defined as chasing, sniffing, or grooming the juvenile conspecific intruder).

#### Social interaction test

An open arena (24” × 24”) with black walls was used for the social interaction test. Two inverted wire cups of diameter 4” and height 4.25” (Spectrum Diversified Galaxy Pencil Holder) were placed in the arena in opposite corners, each set 6” away from the corner of the arena. Red lab tape was placed on the floor of the arena around the outside of the wire cup to demarcate a diameter 4 cm larger than that of the cup. A juvenile male conspecific (PD 24-28) was placed into one wire cup (herein referred to as conspecific cup), while the other cup was left empty (herein referred to as empty cup). The test animal was placed into the middle of the arena and was allowed to freely explore the arena for 10 minutes. Experimenters scored the total amount of time that the experimental animal spent within each region outlined by tape, and computed the time spent with the conspecific cup, as well as the difference score (percent time spent with empty cup subtracted from percent time spent with conspecific cup).

#### Female exposure

One female mouse (PD 30-40) was placed into a clean homecage with a clear acrylic, ventilated cage top, which was used as the interaction chamber. The experimental male mouse was then placed into the chamber and was allowed to interact freely for 5 minutes. The amount of time the male mouse interacted with the female - defined as sniffing, chasing, mounting, or other contact initiated by the male - was manually scored.

### 2.7 Neurogenesis

In a separate cohort of animals, stereotaxic surgery was performed to infuse pAAV_9_-*c-Fos-*tTA + pAAV_9_-TRE-ChR2-eYFP into the dDG and mice were then left undisturbed to recover for 10 days. On day 11, animals were taken off dox for 48 hours and left undisturbed in their homecages. Animals were then split into 3 groups: footshock (negative), novel homecage (neutral), or female exposure (positive) (refer to: *Experience tag)*. Cells active during these behavioral epochs were labelled. Mice were then subjected to the chronic optogenetic stimulation protocol (see *Chronic Stimulation Protocol*) and were then left undisturbed in their homecages for 7 days to allow for optimal doublecortin expression (Couillard-Despres et al., 2005). On the 8th day, animals were euthanized, and their brains were extracted for immunohistochemical staining (see *Immunohistochemistry)*. Doublecortin-positive cells in the upper and lower blade of the DG granule cell layer were manually counted by an experimenter (see *Cell Counting*)

### 2.8 Immunohistochemistry

Mice were overdosed with isoflurane and perfused transcardially first with 40 mL ice cold 1X phosphate-buffered saline (PBS) followed by 40 mL ice cold 4% paraformaldehyde in PBS. Brains were extracted and stored at 4°C, first in 4% paraformaldehyde for 48 hours, and subsequently in PBS. A vibratome was used to obtain 50-μm coronal slices, which were stored in 24-well plates in PBS at 4°C. These slices were incubated with 1X PBS with 2% Triton (PBS-T) + 5% normal goat serum (NGS) for one hour at room temperature for blocking. Primary antibodies were diluted in PBS-T + 5% NGS as follows: guinea pig anti-c-Fos (1:1000, Synaptic Systems, #226 004), chicken anti-GFP (1:1000, Invitrogen, #A10262), and rabbit anti-doublecortin (1:500, Synaptic Systems, # 326 003). Slices were incubated in the primary solution at 4°C for 24 h on an orbital shaker. This was followed by three 10 minute washes in PBS-T, shaking at room temperature. Slices were then incubated with a secondary antibody solution for 2 h at room temperature, shaking. Secondary antibodies were diluted in PBS-T + 5% NGS as follows: Alexa 555 goat anti-guinea pig (1:200, Invitrogen, #A21435), Alexa 488 goat anti-chicken (1:200, Invitrogen, #A11039), and Alexa 555 goat anti-rabbit (1:200, Invitrogen, #A21429). Again, this was followed by three 10-minute washes in PBS-T at room temperature, shaking. Slices were then mounted onto microscope slides with VECTASHIELD® Hardset™ Antifade Mounting Medium with DAPI (Vector Labs, #H-1500).

### 2.9 c-Fos quantification

The total number of neurons immunoreactive for c-Fos were counted in several brain regions - prefrontal cortex (PFC), nucleus accumbens core (NAcc Core), nucleus accumbens shell (NAcc Shell), lateral septum (LS), dorsomedial hypothalamus (dmHyp), lateral hypothalamus (LatHyp), dorsal CA1 (dCA1), dorsal CA3 (dCA3), basolateral amygdala (BLA), and lateral habenula (LHb) - to measure neuronal activity in these areas during defined behavioral assays (female exposure test, resident intruder test, and social interaction test). Animals were euthanized 90 minutes following these tasks, to maximize the robustness of c-Fos expression. Following brain extraction, 3 coronal slices were selected from each of the following regions: approximately +1.15 AP, −2.2 AP, and −2.78 AP. This allowed for visualization of the six brain regions of interest. Following c-Fos staining, the brain regions of interest were imaged using a confocal microscope (Zeiss LSM-800). Images were then processed using FIJI software. The Despeckle tool was used to reduce background noise, and the Subtract Background tool was used to create greater contrast between cells and background. Each brain region z-stack was set to include a 320 μm × 320 μm region of interest (ROI), then processed using the 3D Iterative Thresholding of the 3D ImageJ Suite (Ollion, Cochennec, Loll, Escudé, & Boudier, 2013). The settings for thresholding were set constant for each brain region of each cohort, with a minimum threshold and preliminary size filter, to maintain consistency in image processing and cell counting parameters between animals. The thresholded images were then z-projected to create flat images of the thresholded objects discovered by the plug-in. In order to isolate cell objects from artifacts such as blood vessels or noise, the images were then run through a pipeline created in Cell Profiler 3.1.8 software that identified objects of a particular (more stringent) size and shape. The number of c-Fos-positive cells was recorded for each ROI and averaged within each animal.

### 2.10 Doublecortin quantification

Following chronic reactivation of dDG cells encoding a negative, neutral, or positive experience, animals were left undisturbed in the homecage for 7 days to allow for optimal expression of doublecortin, a marker of neurogenesis. The number of neurons in dDG and vDG immunoreactive for doublecortin (DCX) was examined to determine levels of neurogenesis. In FIJI software, DCX-positive cells were selected in each layer in the z-stack with the Oval tool and added to the ROI Manager. Only DCX-positive cells in a 600 μm × 100 μm ROI were counted. Cells in the dorsal and ventral blades of the dentate gyrus were counted separately.

### 2.11 dDG target verification and ensemble size quantification

In all cohorts, immunoreactivity for eYFP (by proxy of anti-GFP staining) was examined to ensure bilateral expression of the virus in targeted regions. Animals that did not show eYFP immunoreactivity bilaterally in the target region were excluded from analysis. Activity-dependent ensemble size was determined using a subset of animals in each group; the number of eYFP-positive cells in a 600 μm × 100 μm ROI in the dorsal blade of dDG was manually quantified using FIJI software as described above.

### 2.12 Statistical Methods

Calculated statistics are presented as means +/- SEM. To analyze differences, we used two-way repeated measures (RM) ANOVAs (between subject factor: Group; within-subject factor: Time). When time was not a factor, we used one-way ANOVAs. When appropriate, these tests were followed up with post-hoc analyses (Tukey HSD; Sidak, and *a priori* t-tests). All data were tested for normality using Shapiro-Wilk test and homogeneity of variance was assessed with Levene’s test. In the case of the necessity of non-parametric statistics, Kruskal Wallis tests were used. Data were analyzed using GraphPad Prism 8.0 and SPSS Statistics v26 software. Alpha was set to 0.05. All tests were two-tailed.

### 2.13 Data Availability

All relevant data supporting the findings of this study are available from the corresponding author upon reasonable request.

## 3. Results

Hippocampal cells were tagged during either a positive, negative, or neutral behavioral epoch, a design that was implemented to allow for stimulation of similarly sized cellular ensembles encoding experiences of different valences. Mice showed no differences across groups in terms of the number of eYFP cells labelled in the dDG (One-Way ANOVA, F_2,13_ = 1.392, p = 0.2834) (Figure 1C-D). Animals then underwent a previously established (Chen et al., 2019; Ramirez et al., 2015) 10 minute optogenetic stimulation protocol twice daily for 5 days, followed by a behavioral test assessing social behaviors (Figure 1E).

To test the effects of chronic stimulation on social behaviors, male mice underwent a female exposure test after chronic stimulation (Felix-Ortiz, Burgos-Robles, Bhagat, Leppla, & Tye, 2016) (Figure 1F). A Two-Way RM ANOVA revealed that mice did not differ in their baseline levels of interaction with a female as there was no main effect of group (F_2,21_ = 0.0303, p = 0.9702). However, there was a main effect of time (F_1, 21_ = 22.30, p = 0.0001) when compared to pre-stimulation baseline such that male mice interacted more in the post-stimulation test (Figure 1F). While there was a general increase over time in all groups, post hoc analyses revealed that this effect was driven by differences in the positive (p = 0.0128) and negative (p = 0.0073) groups suggesting that chronic stimulation of a salient or valent experience can increase the propensity to interact socially and this may be more pronounced after stimulation of an aversive cellular ensemble in particular. Similar to baseline, there were no group differences during the post-stimulation female exposure test.

We next assessed if chronically reactivating hippocampus-mediated memories affected social behavior involving only males using two additional tests: social interaction and resident intruder (Figure 1G-H). Surprisingly, chronic optogenetic stimulation of dDG cells involved in the encoding of a positive, negative, or neutral behavioral epoch did not result in group differences in time spent interacting with a novel, juvenile conspecific male in the social interaction test (One-Way ANOVA for time interacting F_2,25_ = 0.09415, p = 0.9105 and difference score F_2,25_ = 0.1382, p = 0.8716) or the resident intruder test (One-Way ANOVA F_2,23_ = 0.3150, p = 0.7329). Notably, as mice were not administered a baseline test of these measures prior to chronic stimulation due to the nature of our “tagged” experience, these comparisons could only be made at the post stimulation time point, therefore within-group changes in social interaction or resident intruder behaviors could not be assessed.

Given the observed within animal differences in time spent interacting with a female mouse pre-vs. post-stimulation, we sought to determine whether stimulation of ensembles encoding negative, neutral or positive memories had lasting effects on regional brain activity. Previous research has shown that brain-wide expression of c-Fos is distinct during male or female social behaviors (Kim et al., 2015). Therefore, we quantified the mean number of c-Fos+ cells in various brain regions implicated in processing social interaction and valence (Figure 2B). We found no group differences in the mean number of c-Fos+ cells per area in all brain regions observed (see figure legend for statistics in each brain region). Surprisingly, neurogenesis in the DG was not differentially affected by chronic reactivation of dDG neuronal ensembles. Previous studies have found that chronic optogenetic stimulation of cells encoding female exposure rescues stress-induced deficits in neurogenesis in the DG and social behaviors are known to modulate levels of neurogenesis (Gheusi, Ortega-Perez, Murray, & Lledo, 2009; Opendak, Briones, & Gould, 2016; Ramirez et al., 2015). However, in the current study, we quantified cells expressing doublecortin, a neuronal marker for immature neurons, and found no group differences in expression (Figure 3B-E). Chronically stimulating positive, neutral, or negative dDG ensembles had no effect on neurogenesis in the dorsal (One Way ANOVA, F_2,10_ = 0.4617, p = 0.6430) or ventral (F_2,10_ = 0.1272, p = 0.8819) DG, suggesting that the observed effects of chronic optogenetic stimulation of differentially-valent experiences on female interaction were not likely due to underlying changes in neurogenesis. However, the lack of long-term effects on c-Fos and neurogenesis aligns with lack of group differences in post-stimulation behaviors.

**Figure 2.**
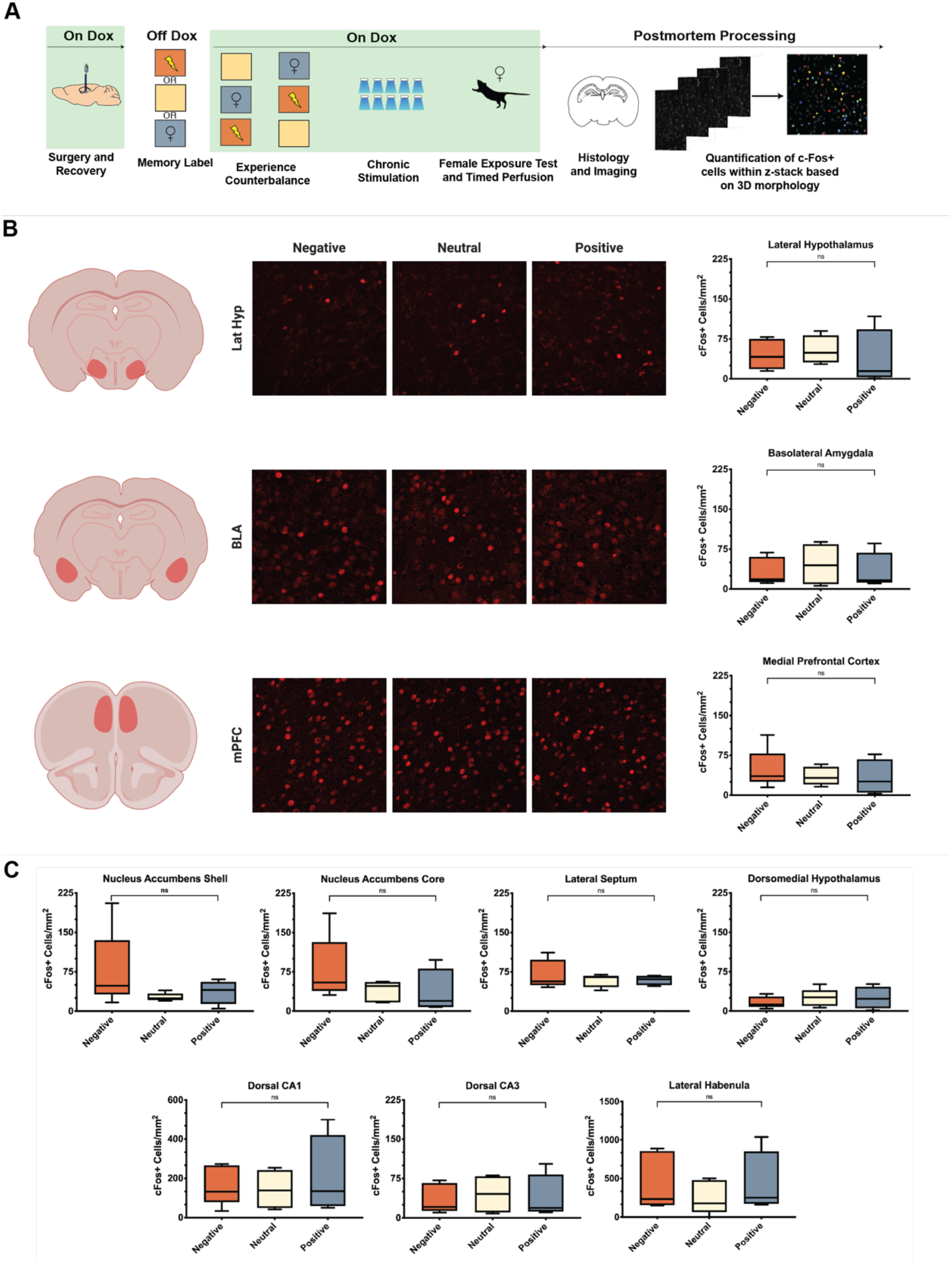
Chronically stimulating dDG ensembles encoding a foot shock, novel homecage, or female exposure, experience does not differentially affect c-Fos across multiple brain regions. (**a**) Behavioral schedule and groups used to examine brainwide c-Fos activation during female exposure after chronic stimulation of different dDG ensembles. Green regions depict periods in which dox was present in the diet, and white regions depict regions where dox was removed to tag active cells (“memory label”). The orange box with a shock symbol represents a four-shock protocol (Negative, n=5), the tan-colored box represents a clean homecage exposure (Neutral, n=5) and the gray box with a female symbol represents exposure to a female conspecific (Positive, n=4). (**b**) Representative images depicting expression in the lateral hypothalamus (Lat Hyp), basolateral amygdala (BLA) and medial prefrontal cortex (mPFC) and quantification of c-Fos activation during female exposure after chronic stimulation of dDG negative, neutral and positive ensembles. (One-Way ANOVAs, lat Hyp: F_2,10_ = 0.2025, p=0.8200; BLA: F_2,10_ = 0.2163, p = 0.8091; mPFC: F_2,11_ = 0.3545, p = 0.7093) (**c**) Mean c-Fos+ per area during post-stimulation female exposure for nucleus accumbens core and shell (NAc Core, One-Way ANOVA: F_2,11_ = 1.356, p = 0.2976; NAc Shell, One-Way ANOVA: F_2,11_ = 1.581, p=0.2492, Lateral Septum (LS, Kruskal-Wallis test, H = 0.28, p = 0.8791), Dorsomedial hypothalamus (dmHyp, One-Way ANOVA: F_2,11_ = 0.4055, p = 0.6762), dorsal CA3 (dCA3, One-Way ANOVA: F_2,10_ = 0.07539, p = 0.9279), dorsal CA1 (dCA1, One-Way ANOVA: F_2,10_ = 0.2, p=0.8220) and lateral habenula (LHb One-Way ANOVA F_2,11_ = 0.5030, p = 0.6180).

**Figure 3.**
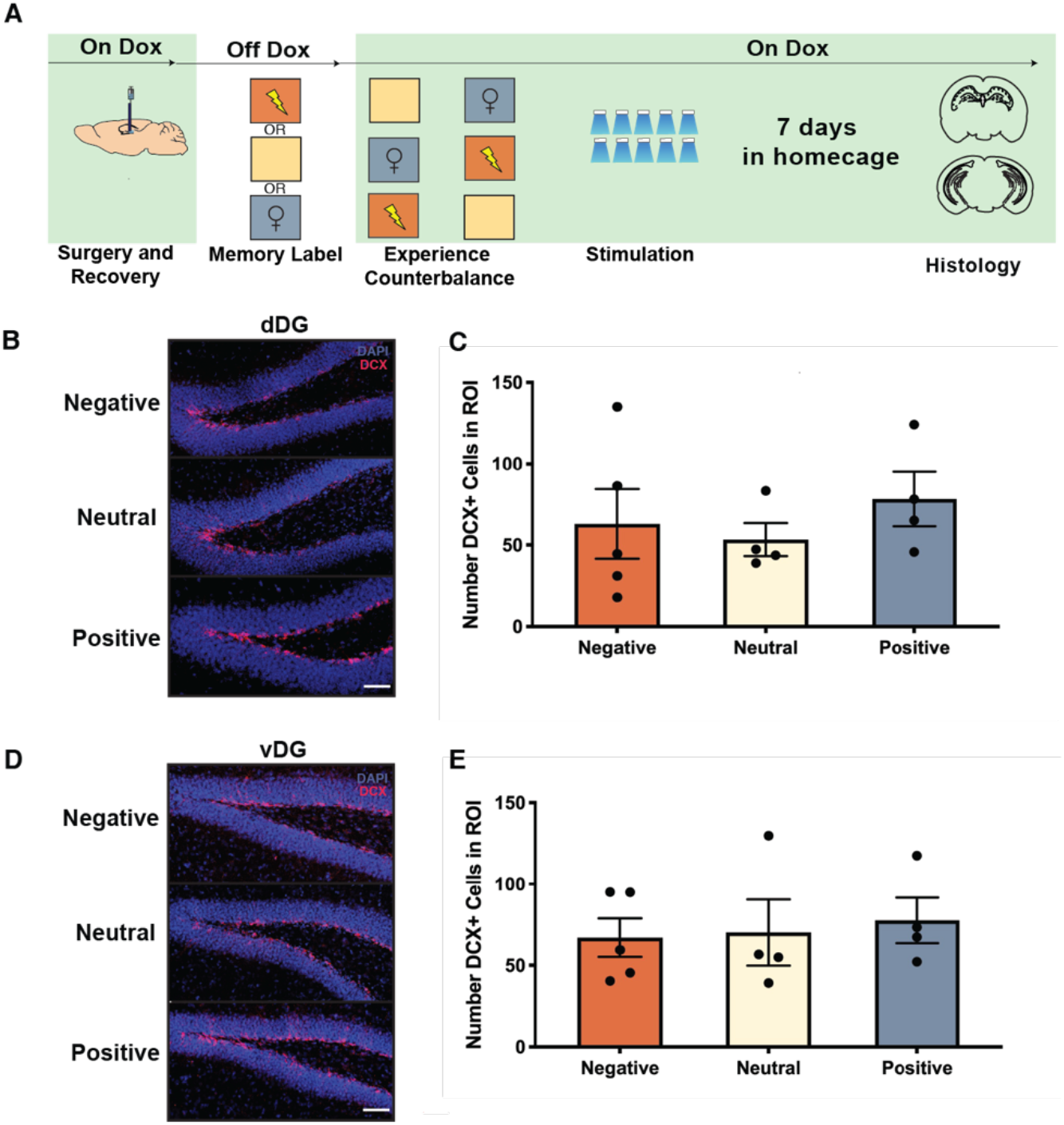
Chronically stimulating dDG ensembles encoding a foot shock, novel homecage, or female exposure, experience does not alter neurogenesis in dDG or vDG. (**a**) Behavioral schedule and groups used to examine neurogenesis induced by the chronic stimulation protocol. Green regions depict periods in which dox was present in the diet, and white regions depict regions where dox was removed to tag active cells (“memory label”). The orange box with a shock symbol represents a four-shock protocol (Negative, n=5), the peach-colored box represents a clean homecage exposure (Neutral, n=4) and the gray box with a female symbol represents exposure to a female conspecific (Positive, n=4). (**b**) Representative images of dDG (**c**) and quantification of doublecortin-positive cells (red) in the dDG for each group (One-Way ANOVA, F_2,10_ = 0.4617, p = 0.6430). (**d**) Representative images of vDG and quantification (**e**) of doublecortin-positive cells (red) in the vDG for each group (One-Way ANOVA, F_2,10_= 0.1272, p=0.8819). Data are presented as mean ± s.e.m. Scale bar represents 100 μm. Dorsal dentate gyrus (dDG), ventral dentate gyrus (vDG).

## 4. Discussion

Our findings demonstrate that chronic stimulation of dDG neurons involved in the encoding of a salient behavioral experience can drive an increase in male-female interactions post-stimulation, an effect partially modulated by the valence of memory stimulated, consistent with the hippocampus’ role in processing both mnemonic and valence-related information (Fanselow & Dong, 2010). Interestingly, stimulation of cells encoding an aversive (footshock) or socially appetitive (female encounter) experience both drove enhancement of subsequent female interaction. Thus, reactivating cells that encoded experiences of opposite valence modulated behavior similarly, when comparing footshock and female exposure groups to mice that experienced stimulation of cells encoding a novel homecage exposure. While our results suggest a mild increase in subsequent female interaction within the neutral group, this increase failed to reach statistical significance, potentially indicating that while cells processing a novel homecage exploration are sufficient to act as a functional conditioned stimulus when acutely activated (Ramirez et al., 2013), they may not be sufficient to drive differences in social behaviors when chronically stimulated. Together, this suggests stimulation of cells that process more salient experiences, such as footshock or female exposure, is more effective for driving changes in subsequent female interaction.

We propose that these findings add a valence- and experience-specific social element to previous studies that measured hippocampus-mediated memory recall in which stimulation of differentially valent memories drove bidirectional effects on behavioral responses (Chen et al., 2019; Ramirez et al., 2015). While these studies found that chronic stimulation induced behavioral changes specifically at post-stimulation timepoints, we speculate that our lack of post-stimulation behavioral and regional brain activity *across* groups are a result of varying stimulation protocols (chronic vs. acute) and timing of stimulation (i.e. during a behavioral testing session) within an experimental protocol. For instance, chronic stimulation may affect behaviors between groups only following chronic stress, and acute optogenetic stimulation may alter behavior during a stimulation session but may not be sufficient to induce lasting structural or functional changes supporting enduring behavioral effects (Ramirez et al., 2015; Redondo et al., 2014). Overall, these findings suggest chronic stimulation of distinct hippocampal memory ensembles may drive variable behavioral responses depending on the specific cellular ensemble reactivated and type of behavior measured.

Moreover, as reactivation of dDG cells encoding both a fearful experience or a novel female encounter drove a subsequent increase in interaction with a female, we speculate that continued reactivation of a positive memory engram cells may reinforce the downstream responses promoting female interaction or mate-seeking behavior, in a manner perhaps similar to how chronic stimulation of a contextual memory can bi-directionally modulate the original memory itself (Chen et al. 2019). However, it is less clear why continued reactivation of cells encoding a fear memory would also increase subsequent female interaction. We posit that chronic stimulation extinguished fear responses and thereby encouraged female interaction, as chronic reactivation of dDG fear memory ensembles has been shown to reduce contextual fear responses (Chen et al. 2019). Alternatively, it is possible that repeated reactivation of dDG cells encoding fear caused mice to seek female interaction further, as female interaction has also been shown to attenuate fear (Bai, Cao, Liu, Xu, & Luo, 2009). It is important to note that with our experimental design, it is possible that the within-group enhancement of female interaction time was driven by a combination of chronic optogenetic stimulation of dDG populations encoding different valenced experiences and by different orders of experienced behaviors, though we believe this is unlikely as we observed no group differences in the pre-stimulation female interaction test. Finally, a second female encounter may contribute to this enhancement of female interaction across time, as well. However, due to the stronger effect observed in the negative (shock) and positive (female) stimulation groups relative to chronic stimulation of a dDG population encoding a neutral (homecage) experience, we believe that the stimulation of the negatively and positively valenced dDG neuronal ensembles further enhanced female interaction beyond the effects of re-exposure to a female per se.

Interestingly, we did not find group differences in social behavior when comparing groups that had received chronic stimulation of negative, neutral, or positive dDG populations during the post-stimulation behavioral test (Fig 1F-H). This suggests that while chronic stimulation of dDG ensembles encoding fear or female exposure may facilitate subsequent female interaction in a within-subject manner across time, chronic stimulation of these different ensembles does not result in between-group differences at a single time point, possibly due to the differences in optogenetic protocols discussed above. These data are supported by our observations that chronic stimulation of dDG memory ensembles did not yield group differences in c-Fos expression throughout several brain regions, or result in changes in neurogenesis in the ventral or dorsal DG. While our design did not implement a pre-stimulation test for social interaction or resident intruder behaviors, future studies can assess if performing these tests would result in a similar within-group increase in social interaction with our chronic stimulation protocol.

Our results shed further light on the relationship between optogenetic stimulation and neurogenesis in the DG. In our previous reports, chronic activation of a positive memory reversed the effects of stress on neurogenesis, highlighting putative differential effects of stimulating hippocampal cells in a stressed or unstressed rodent (Ramirez et al., 2015). Our results indicate that chronic stimulation of dDG ensembles alone is not sufficient to do so, suggesting that changes in neurogenesis induced by chronic optogenetic stimulation may occur only when administered after stress. It is possible that stress reduces cognitive flexibility, which chronic optogenetic stimulation is sufficient to circumvent and that such perturbations in a healthy rodent have already reached a “ceiling effect” in their capacity to modulate the production of adult-born cells (Anacker & Hen, 2017). Our recent work suggested a bi-directional role for the dorsal and ventral hippocampus in respectively suppressing or enhancing context-specific memories, and we speculate that the capacity for chronic stimulation of hippocampal cells to alter neurogenesis levels too may depend on the site stimulated (Anacker et al., 2018; Chen et al., 2019). Importantly, our findings indicate that chronic stimulation of sparse DG populations does not have adverse, off-target effects on neurogenesis, which is a promising assurance for future studies employing similar chronic optogenetic stimulation in the same brain region.

Together, our results point to the need for future research aimed at understanding the varying effects of chronic stimulation on different brain areas or specific sets of cells stimulated. For instance, stimulation of differently valenced dDG ensembles failed to differentially affect social behavior across groups during post-stimulation social interactions, and this may be due to dorso-ventral differences in the encoding of contextual, emotional or social information, which underscores the ventral DG’s prominence in processing similar types of information (Ciocchi, Passecker, Malagon-Vina, Mikus, & Klausberger, 2015; Kheirbek et al., 2013; Okuyama, Kitamura, Roy, Itohara, & Tonegawa, 2016), its influence on neurogenesis (Anacker et al., 2018), and its putative promise as a future target for chronic stimulation.

Finally, a myriad of recent studies have leveraged the effects of repeatedly activating various brain regions and circuits to note their enduring effects on behavior. For instance, optogenetic-induced long-term depotentiation was sufficient to lastingly impair a memory while subsequent induction of long-term potentiation restored the expression of the memory (Nabavi et al., 2014). Additionally, high-frequency spike trains that lasted for 10 minutes were sufficient to alter excitation/inhibition balance and spine levels in the hippocampus and also facilitated the extinction of a contextual memory (Mendez, Stefanelli, Flores, Muller, & Lüscher, 2018), which points to the power of prolonged optogenetic strategies in modifying the structural and functional properties of the hippocampus. Various groups have also utilized optogenetic-inspired deep-brain stimulation strategies to provide a translational approach to enduringly reprogram a brain out of a maladaptive state (Creed et al., 2019), and we propose that artificially manipulating engrams provides a conceptual means by which to resculpt neural activity and behavior.

Overall, our data suggest that chronic stimulation of hippocampus-mediated memory engrams can differentially affect social behaviors over time without inducing widespread changes in c-Fos or neurogenesis, reinforcing the importance of considering multiple factors such as off-target effects (Otchy et al., 2015), the specific behavioral assays used, and which measures of neural changes will be analyzed when implementing chronic stimulation protocols.

## Acknowledgements

We would like to thank Joshua Sanes and his lab (Center for Brain Science, Harvard University) for providing laboratory space within which the initial experiments were conducted; Harvard University’s Center for Brain Science Neuroengineering core for providing technical support. We thank Abby Finkelstein for helpful comments on the manuscript. Finally, we would like to thank Susumu Tonegawa and his lab (Massachusetts Institute of Technology) for providing the activity-dependent virus cocktail. This work was supported by an NIH Early Independence Award (DP5 OD023106-01), a Transformative R01, Young Investigator Grant from the Brain and Behavior Research Foundation, a Milton Grant from the Society of Fellows at Harvard University, a Ludwig Family Foundation Grant, and the McKnight Foundation Memory and Cognitive Disorders Award.

## Author Contributions

E.D., H.L., A.M., C.C., J.L., S.L.G., and S.R. designed and performed the experiments. E.D., A.M., H.L., and S.R. wrote the paper. All authors edited and commented on the manuscript.

## Competing Financial Interests

The authors declare no competing interests.

## Notes

### Competing Interest Statement

The authors have declared no competing interest.

